# Growth characteristics of natural microbial populations are skewed towards bacteria with low specific growth rates

**DOI:** 10.1101/2024.04.24.590925

**Authors:** Ashley N Bulseco, Wenzhou Yang, Julie A Huber, Joseph J Vallino

## Abstract

Microbes are often functionally characterized by traits that specify their optimum environmental conditions for growth, such as temperature or pH, as well as upper and lower bounds where growth is possible. While any given microbe will have a narrow environmental window where growth can occur, a diverse community can span a much larger range of conditions where growth is possible by at least some members of the community. One important trait of microbes is maximum specific growth rate, as this trait determines if a microbe can persist in environments with short residence times. In this study, we conducted a chemostat experiment with natural microbial communities and manipulated dilution rate to test how it would act as a selective force controlling community dynamics and microbial diversity. Using both experimental chemostats and trait-based modeling, we examine how the composition of a microbial community collected from a coastal meromictic pond changes as dilution rate increases from 0.1 to 10 d^−1^. We compare experimental results from 16S rRNA gene amplicon sequences to the output from two different simulations of the trait-based model, one in which maximum specific growth rate is initially evenly distributed across the community and another where maximum specific growth rate traits are pulled from a beta probability distribution that is skewed towards low specific growth rates. Our experimental results match the simulation where the initial natural population is dominated by slow-growing microbes. Our results also highlight the importance of initial trait distributions in modeling community response to environmental changes.

**Importance:** All living organisms are constrained by environmental boundaries that govern where growth is possible, such as minimum and maximum temperature or pH. An organism’s maximum specific growth rate places a lower bound on the residence or turnover time of a system where an organism can persist without being removed from the system. While we have good bounds on the maximum specific growth rate of culturable organisms, such as *Escherichia* or *Vibrio* species, information is lacking on how the maximum specific growth rate trait is distributed in natural microbial communities. This information is critical for understanding how a community will respond to changes in residence time. Using chemostats and trait-based modeling, this study assesses how maximum specific growth rate is distributed in a natural community collected from a coastal, salinity-stratified pond.

## Introduction

Microbes are among the most abundant, diverse, and ubiquitous organisms on Earth. As such, their interactions mediate important biogeochemical cycles, influencing ecosystem processes such as nutrient fluxes, productivity, and carbon cycling (Falkowski et al. 2008). A major goal of microbial ecology is to understand and predict what ecological mechanisms drive biodiversity, structure, and dynamics in microbial communities, as these characteristics are linked to ecosystem function and biogeochemistry. However, characterizing these mechanisms continues to be a challenge given the high diversity of microbes (Sogin et al. 2006) and the proportion of microbes across environments that remain uncultured (Steen et al. 2019). Modeling can begin to overcome some of these challenges (Follows & Dutkiewicz 2011), though models based on explicit representation of all microbial species present in a system are problematic. Trait-based modeling approaches aim to identify and improve our understanding of mechanisms underlying ecological patterns that structure microbial communities through the exploration of functional traits, which relaxes the emphasis on individuals (Follows et al. 2007, Green et al. 2008, Litchman et al. 2015, Lajoie & Kembel 2019). These traits, which can be morphological, physiological, or genomic, incorporate physiological trade-offs and thus constrain the performance of an organism under a set of given environmental conditions (Violle et al. 2007, Krause et al. 2014). Organisms with particular functional traits that are better suited for a given environment are favored, and this environmental selection ultimately determines the structure of the microbial community and the dominant ecosystem processes (Wallenstein & Hall 2012, Malik et al. 2020), as well as how these linkages may change in the future, provided tradeoffs between traits are properly understood (Litchman et al. 2015).

Microbial life in extreme habitats serves as one example where the external environment selects for organisms with specific functional traits (Rothschild & Mancinelli 2001). As habitats approach extremes in temperature, salinity, pH, or other boundaries, the diversity of microbial communities decreases and becomes dominated by only a few taxa best adapted to the given specialized conditions (Lewin et al. 2013). For instance, Sharp et al (2014) found a strong unimodal decrease in microbial diversity (Shannon index) at temperature extremes, and a similar, but weaker, relationship with pH extremes from terrestrial springs. In arable soil, Rousk et al. (2010) found a strong, positive relationship between bacterial diversity and pH, with effectively half the number of operational taxonomic units (OTUs) existing at pH 4 compared to a pH of 8.3. Similar patterns of low diversity have been described in other extreme temperature environments, such as hot springs in Yellowstone (Bhaya 2012, Inskeep et al. 2013) and hydrothermal vent systems (Frank 2013), as well as acidic mine-drainage habitats (Baker & Banfield 2003, Tringe et al. 2005, Teng et al. 2017). In each of these examples, diversity decreases as the environmental boundaries of life that define the habitable zone are approached, because fewer organisms are capable of surviving in niches defined near a boundary. Where boundaries are crossed, life is excluded.

Resource supply rate and associated transport processes is another type of extreme that planktonic organisms must also contend with. In systems subject to advective and diffusive transport (Vallino & Hopkinson 1998), such as streams, lakes and meso-scale eddies, turnover rate is defined by the volumetric rate of water entering and leaving the system (e.g., m^3^ d^−1^) divided by the volume of the system (m^3^), and residence time is its inverse. For microbes, the specific growth rate is defined by growth rate (cells L^−1^ d^−1^) divided by their abundance (cells L^−1^). Consequently, an organism’s specific growth rate must match or exceed the water’s turnover rate; otherwise, it will be washed out of the system regardless of whether or not it is a good competitor. Since specific growth rate depends on resource concentration (Holling 1965), an organism can persist in a system with high water turnover rates provided resource concentration is high and water turnover rate does not exceed the organism’s maximum specific growth rate. Maximum specific growth rate and resource affinity are also important in structuring microbial communities, as competitive abilities between organisms depend on both processes and has been widely studied in ecology. For instance, the trade-off between maximum growth rate and resource affinity, the r-K strategy dichotomy (Macarthur & Wilson 1967, Sommer 1981) or the similar R* rule (Tilman 1980), describes a set of metabolic strategies that favor either high resource affinity or high maximum growth rates, but follows a continuum from “gleaners” to “opportunists”, respectively (Fredrickson & Stephanopoulos 1981, Grover 1990, Litchman et al. 2015). Transport processes play an important role in competition as well, and Locey and Lennon (2019) provide a review of theories governing residence time (the inverse of turnover or dilution rate) and biodiversity.

Trait-based models can be used to study competitive interactions between microbes for resources that are subject to transport process, but how traits are initialized and distributed across the community is important to consider as the communities approach conditions near growth extremes that are a function of maximum specific growth rate and substrate affinity. For instance, for a natural microbial community, we can define a trait variable that governs the tradeoff between high specific growth rate and high substrate affinity that represents a continuum from oligotrophs using ATP-binding cassette transporters at one end of the spectrum, to copiotrophs using phosphotransferase systems (Norris et al. 2020) at the other end. How is this trait distributed in natural communities with high richness? Are copiotrophs just as likely as oligotrophs, or is the community skewed towards one side or the other? The distribution of this trait should impact community composition as the growth rate boundary is approached and selects for microbes with high specific growth rates.

To test how growth rate and substrate affinity traits might act as a selective force controlling microbial community dynamics and diversity, we conducted a chemostat experiment with manipulated dilution rate using a natural microbial population collected from a coastal pond. The use of a chemostat experiment is advantageous because the convergence of cellular growth and dilution rate at steady state allows for precise characterization of microbial dynamics under a given environmental condition (Smith & Waltman 1995), and the specific growth rate of microbes must be able to match or exceed the chemostat’s dilution rate; otherwise, they will be washed out. Here, we hypothesize that as dilution rate is increased, the diversity of the community will decrease because natural communities are skewed towards oligotrophic strategies. In contrast, if the pond microbial community has a uniform distribution in specific growth rate vs substrate affinity trait, then diversity will not change as dilution rate increases, at least prior to complete washout. To test our hypothesis, we developed a simple trait-based model that predicts changes in biodiversity assuming either 1) a uniform probability distribution of bacterial growth kinetic traits (null hypothesis) or 2) a beta probability distribution of traits favoring lower specific growth rates, mimicking conditions we hypothesized to be present in nature. By comparing empirical observations with the trait-based model simulation, we find that diversity diminishes as the growth rate boundary is approached in both experimental and model results using the skewed distribution, supporting the hypothesis that the pond microbial community diversity is weighted towards oligotrophs (aka, gleaners). Our results also demonstrate the importance of trait initialization on the performance of trait-based models.

## Methods

### Experimental Setup

We collected a 10 L water sample at 2.5 meters below the surface from a coastal, meromictic pond (Sider’s Pond in Falmouth, MA; 41°32’52.08” N, −70°37’26.82” W), which is permanently stratified and characterized as both eutrophic (Caraco 1986) and phosphorus limited (Caraco et al. 1987). *In situ* water conditions were 12.8°C with salinity and pH of 5.48 and 7.09, respectively. After passing water through a 300 *μ*m Nytex mesh to remove large organisms and particles, we inoculated two replicate (MC1 and MC2) 3 L bioreactors (Bellco glass: 1964-03600, outfitted with PTFE low shear impellers 1964-03404) with 3 L of Siders Pond water. We chose to start with 100% pond water, rather than inoculating a defined media to maximize inclusion of the rare biosphere (Sogin et al. 2006) and functional diversity (Lynch & Neufeld 2015).

Environmental operating conditions were maintained at 25°C in the dark by a environmental chamber (PGR15 Conviron, Winnipeg, Canada), and chemostats were sparged with synthetic air from a gas cylinder (21.06% O_2_, 0.04% CO_2_, balance N_2_) at a flow rate of 10 sccm (10.92 mL min^−1^ at 25°C and 101.3 kPa) regulated with mass flow controllers (MKS M100B, Andover, MA). To prevent growth in the medium feeding the chemostats, it was divided into two parts consisting of a nutrient only and carbon only media. The nutrient medium (NM) consisted of 15 *μ*M KNO_3_, 2 *μ*M HK_2_PO_4_, 100 *μ*M MgSO_4_ 7H_2_O, 100 *μ*M CaCl_2_ 2H_2_O, 1 mL of a concentrated trace element (TE) solution per L of NM (Fernandez-Gonzalez et al. 2016), and adjusted to a salinity of 3 PSU using Instant Ocean Sea Salt. The TE solution contained 18.5 mM FeCl_3_ 6H_2_O, 485 *μ*M H_3_BO_3_, 126 *μ*M CoCl_2_ 6H_2_O, 100 *μ*M CuSO_4_ 5H_2_O, 348 *μ*M ZnSO_4_ H_2_O, 163 *μ*M MnSO_4_ H_2_O, 124 *μ*M Na_2_MoO_4_ 2H_2_O, 84 *μ*M NiCl_2_ 6H_2_O and 10 mL L^−1^ 37% HCl. The carbon medium (CM) consisted of 5.0 mM glucose, 5.969 mM xylose, 10.83 mM ethanol, 16.75 mM acetate, and 20.61 mM methanol, where carbon substrate concentrations were chosen such that each substrate contributed 14.7 kJ L^−1^ Gibb’s free energy at 293K and pH 6.8 to the total (Alberty 2003). Dedicated pumps (Masterflex 07523-90 drive, Easy-Load 3 pump head, and L/S Norprene tubing) were used for the NM and CM, and the CM pump was also fitted with two additional pump heads (4 in total) with larger L/S tubing, which were used to withdraw media from the chemostats to maintain constant bioreactor volume. To manipulate resource availability, we supplied the nutrient plus carbon media at three different dilution rates and two different media ratios (CM:NM) as follows: 0.1 d^−1^ (0.3 L d^−1^) with a 1:98.3 CM:NM ratio; 1.11 d^−1^ (3.33 L d^−1^) with a 1:8.24 CM:NM ratio; 10 d^−1^ (30 L d^−1^) with a 1:10 CM:NM ratio. The experiment started at the 0.1 d^−1^ dilution rate and was stepped up to 1.0 and 10. d^−1^ sequentially, which were run for 15.5 d, 5.3 d, and 3 d, respectively (Table 1).

**Table 1.** Experimental chemostat conditions. Samples with asterisks were sequenced successfully in only one of the two chemostat replicates. CM: Carbon medium; NM: Nutrient medium; CM:NM: Ratio between the Carbon and Nutrient medium.

| Dilution Rate (d<sup>-1</sup>) | CM Flow (L d<sup>-1</sup>) | NM Flow (L d<sup>-1</sup>) | CM:NМ | Days Run (d) | Day of Collection |
| --- | --- | --- | --- | --- | --- |
| 0.1 | 0.0030 | 0.30 | 1:98.3 | 15.5 | 0 (pond), 0.8, 5.8*, 9.5*, 10.4, 15.4 |
| 1.1 | 0.36 | 2.97 | 1:8.24 | 5.3 | 16.5, 17.6, 18.8, 19.8 |
| 10 | 2.73 | 27.3 | 1:10.0 | 3 | 21.5, 21.8, 22.4, 22.8*, 23.5 |

### Sample collection & biogeochemical analyses

Every hour, the chemostat monitoring system sampled for gas concentrations of carbon dioxide (CO_2_) and oxygen (O_2_) in the headspace via a closed gas-sampling loop (250 mL min^−1^ total volume) containing a Nafion gas dryer (Perma Pure MD-070-24P, Lakewood, NJ) and an Oxigraf laser diode absorption spectrometer (O2Cap, Sunnyvale, CA). Chemostats were sequentially sampled using a computer controlled Valco (Houston, TX) STF selector valve, which also sampled the feed gas cylinder thereby allowing a single point calibration of the O2Cap gas analyzer to account for instrument drift. We monitored dissolved oxygen (DO) and pH with a Hamilton (Reno, NV) VisiFerm DO Arc 335 and EasyFerm Plus Arc 325 probes, respectively, and controlled and calibrated the probes via Hamilton Device Manager software equipped with a USB-ModBus RS 485 converter.

During each time point (Table 1), we withdrew no more than 500 mL total volume from each chemostat. Using a 60 mL syringe, we filtered sample through a Whatman GF/F filter for nitrate (NO_3_^−^), ammonium (NH_4_^+^), and phosphate (PO_4_^3−^) as well as dissolved organic carbon (DOC), which we preserved with 100 µL of 50% HCl, and stored at −20°C until analysis. For bacterial enumeration, we collected 10 mL of whole water and stored samples at 4°C preserved with 3.7% of formaldehyde (final concentration). For 16S rRNA gene amplicon sequencing, we filtered between 60 and 250 mL of water onto a 0.2 µm polyethersulfone (PES) Sterivex filter (until the filter clogged), which we stored at −80°C until extraction.

We analyzed NO_3_^−^ and PO_4_^3−^ concentrations on a Lachat (QuikChem 8500) and NH_4_^+^ concentrations on a Shimadzu 1601 spectrophotometer using a modified phenol-hypochlorite method (Solorzano 1969). DOC was measured in duplicate on a total organic carbon analyzer (Aurora 1030W, OI Analytical) after sample acidification.

### Bacterial cell counts, library preparation, and sequence analysis

We performed direct bacterial cell counts on water samples. Following methods outlined in Porter & Feig (1980), we filtered 1 mL of sample volume onto a 25 mm, 0.2 µm polycarbonate filter backed by a 0.45 µm GFF filter and fluorescently stained bacterial cells with DAPI (28 µM final concentration, Sigma-Aldrich) for 5 minutes in the dark. We visualized bacterial cells using an epifluorescence Zeiss Axio Imager 2 and performed counts using the 100x objective on 5 fields of view. We extracted genomic DNA from PES filters using the DNeasy PowerSoil Pro kit (Qiagen) following manufacturer’s methods and submitted high quality extracts to the Bay Paul Center at the Marine Biological Laboratory for library preparation and sequencing. Amplicon libraries were prepared using bacterial 16S V4V5 518F (CCAGCAGCYGCGGTAAN) and degenerate 926R primers (CCGTCAATTCNTTTRAGT or CCGTCAATTTCTTTGAGT or CCGTCTATTCCTTTGANT) using Platinum HiFi II DNA Polymerase (Thermo Fisher) and purified using AMPure Beads (Beckman Coulter) at a 0.75x bead-to-template ratio. After quantifying the final pool using a Kapa Biosystems qPCR (Millipore Sigma), sequencing commenced on an Illumina MiSeq, generating 250 nt paired-end reads. These reads can be found in the NCBI Sequence Read Archive under BioProject ID PRJNA1103251.

We used the DADA2 workflow (v1.14.1; Callahan et al. 2016) in R (v3.6.3; R Core Team 2019) to analyze our sequence data. To perform quality filtering, we applied the following parameters in the ‘filterAndTrim’ command: MaxN = 0, maxEE = 2 and 2, truncQ = 2, truncLen = 250 and 225 (based on quality profiles), and trimLeft = 19 and 20, for forward and reverse reads, respectively. After inferring amplicon sequence variants (ASVs) from de-replicated reads, we removed chimeric sequences using the ‘consensus’ method. We assigned taxonomy against the SILVA database (v138; Quast et al. 2013) and filtered out any sequences matching Mitochondria or Chloroplasts due to known primer biases, resulting in 1440 unique ASVs (Table S1). We conducted all downstream sequence analyses on generated ASV tables using the Phyloseq package (v1.30.0; McMurdie & Holmes 2013).

### Statistical analyses

All statistical analyses were conducted in R (v3.6.3; R Core Team 2019). We compared chemostat parameters (*e.g.* %CO2, %O2, DO, and pH) and dissolved nutrients (*e.g.* NO_3_^−^, NH_4_^+^, PO_4_^3−^, and DOC concentrations) across dilution rates using a one-way ANOVA with a Tukey’s HSD test when assumptions of normality and equal variances were met; otherwise, we used a Kruskal-Wallis test by ranks.

To visualize microbial community structure at each sample point, we calculated the relative abundance (%) of ASVs, but we only included those ASVs with abundances greater than 0.1% in stacked bar plots, because we were most interested in large-scale shifts in community structure. We calculated Bray-Curtis dissimilarity on our full dataset with the pond sample removed to quantify diversity between MC samples and visualized the results using a principal coordinates analysis. To estimate dispersion among samples by dilution rate, we calculated the non-Euclidean distance between each sample and group centroids using the ‘betadisper’ function in the vegan package (Oksanen et al. 2019), and compared dispersion among dilution rates using a one-way ANOVA. To better understand microbial community stability between sequential samples, we calculated Bray-Curtis pairwise distance between sequential time points (sampling point 1 compared to 2, 2 to 3, etc). This resulted in a dissimilarity metric, which we then converted to similarity by subtracting 1 from each value such that increasing values indicate greater community stability between samples. That is, a community stability index close to 1 indicates the community composition does not change significantly between two sequential time points. We compared community stability among dilution rates for each MC separately using a one-way ANOVA and identified significantly different means using a Tukey’s HSD multiple comparison test. We then calculated alpha diversity on both rarified and non-rarified datasets using the Shannon index in Phyloseq and compared across dilution rates by MC. Similar analyses also were conducted on simulated ASVs from our trait-based model.

### Trait-based model description

To model the population dynamics of a community of bacteria, we used a previously developed adaptive Monod equation that captures prokaryote growth kinetics across oligotrophic to copiotrophic conditions (Vallino 2011) as the basis for our simulations. Under oligotrophic conditions, bacteria have low specific growth rates (< 1 d^−1^), low growth efficiencies (< 10%), and high substrate affinities (nM substrate concentrations). While under copiotrophic conditions, bacterial specific growth rates can exceed 50 d^−1^, growth efficiencies can exceed 60%, and substrate affinity is low (mM substrate concentrations). In the adaptive Monod kinetics equation, specific substrate uptake rate, *φ*, is governed by a growth efficiency parameter, *ε*, that varies between 0 and 1 to represent growth kinetics across the full range of trophic environments a community may experience, which is given by Eq. (1),

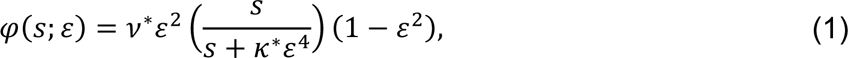

where *s* is substrate concentration (μM), and the community-independent constants *v*^∗^and *k*^∗^ are fixed at values of 350 d^−1^ and 5000 μM, respectively. These two constants have been used successfully for modeling prokaryote growth in both laboratory and field environments (Algar & Vallino 2014, Vallino et al. 2014, Vallino & Huber 2018). To reduce the complexity of the model, the last term in Eq. (1), (1 − *ε*^2^), is an approximate thermodynamic constraint that replaces the formal thermodynamic driver used previously (Vallino & Tsakalakis 2020). The specific growth rate, *μ*, is simply the product of *ε* and *φ*(*s*; *ε*), where bacterial growth is governed by the following stoichiometric equation (2),

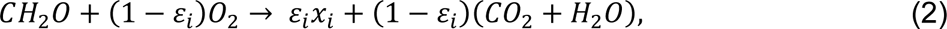

where *x_i_* is bacterial biomass concentration in μM C for instance *i*, which represents a single ASV. For simplicity, bacterial growth is assumed to be solely limited by glucose, *CH*_2_*O*, normalized to unit carbon here. We use trait-based modeling (Follows et al. 2007, Litchman et al. 2015) to simulate the population of bacteria in the experimental chemostats where the growth efficiency, *ε*_*i*_, is a bacterial trait variable that can be assigned a value randomly between 0 and 1. This value represents a continuum of ecological strategies varying from gleaner (*ε*_*i*_ ≈ 0) to opportunist (*ε*_*i*_ ≈ 1). Letting *x_i_*(*t*) be the concentration of a single prokaryote species (or ASV) with trait *ε*_*i*_ at time *t*, the equations governing a population of *n* prokaryotes each with randomly assigned growth efficiency traits in a predator-free chemostat is given by the following 4 equations,

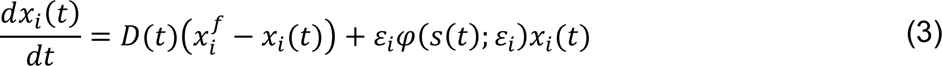

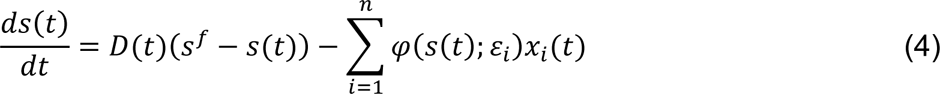

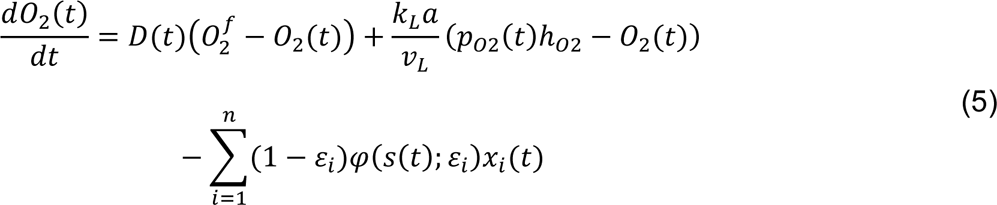

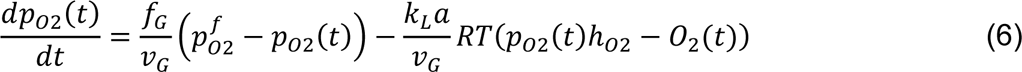

where, *O*_2_ is DO concentration (μM), *p*_*O*2_ is the partial pressure of oxygen in the chemostat headspace (atm), *k*_*L*_ is the liquid-side gas transfer velocity (m d^−1^), *a* is the area of the gas-liquid interface (m^2^), *v*_*L*_ and *v*_*G*_, are volume of the liquid and gas headspaces (m^3^), respectively, ℎ_*O*2_ is the Henry’s law constant for O_2_ in water (μM atm^−1^) (Weiss 1970), *f*_*G*_ is the gas flow rate (m^3^ d^−1^), superscript *f* denotes the state variable concentration or pressure in the input feed, *n* is the number of ASVs in the simulated population, where the values of *ε*_*i*_ are randomly drawn from an appropriate probability distribution, and *D*(*t*) is the dilution rate (d^−1^), set to match the experimental dilution rate over time. We include DO and O_2_ partial pressure in the model to compare with chemostat observations. This allows us to assess the performance of the model relative to the experiment and helps ensure that the model reasonably captures observed process rates. The liquid-side gas transfer velocity, *k*_*L*_, was determined by sparging an operationally sterile reactor with N_2_ until DO was below 25 μM, then rapidly equilibrated the headspace with air and monitored the increase in DO over time while sparing the chemostat with air at the operational flow rate. This process was modeled using Eq. (5), with *p*_*O*2_(*t*) set to atmospheric concentration and *φ*(*s*(*t*); *ε*_*i*_) and *D*(*t*) set to 0. The value of *k*_*L*_ was determined by minimizing the error between predicted and measured DO over time.

The two model scenarios we investigated were, (1) *ε*_*i*_ randomly drawn from a uniform probability distribution, ranging from 0 to 0.7 (Fig. 1A) and (2) *ε*_*i*_ drawn from a beta probability distribution with α = 1.1 β = 13.3 between 0 and 1, which is heavily skewed towards the oligotrophic end of the distribution (Fig. 1B). The first scenario represents a null test of the hypothesis, in that it assumes there is no decrease in the diversity of specific growth rates as the upper limit for prokaryote growth is approached. In contrast, the second scenario assumes there are disproportionately fewer members of the microbial population that have specific growth rates near the maximum possible value of 65 d^−1^ based on Eq. (1). That is, a community skewed towards gleaners.

**Fig. 1.**
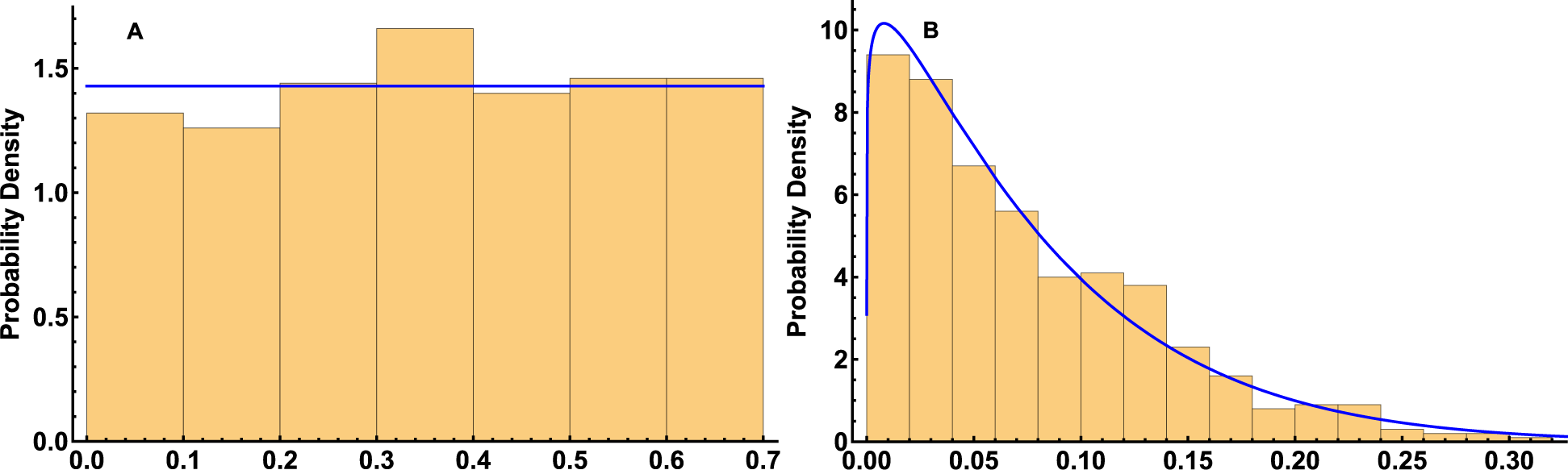
Probability density function (blue lines) and histograms (yellow bars) of 500 samplings of *εi* (growth efficiency of ASV *x_i_*, Eq. (1)) drawn from (A) a uniform distribution over 0 to 0.7 and (B) a beta distribution with α = 1.1 β = 13.3 (B). Note, for the uniform distribution *εi* was limited to 0.7 because the specific uptake rate for the adaptive Monod equation, Eq. (1), begins to decrease for larger values of due to a decrease in the thermodynamic driver. Values of *εi* > 0.7 are theoretically possible, but not evolutionarily competitive. Values of *εi* represents a continuum of ecological strategies varying from gleaner (*εi* ≈ 0) to opportunist (*εi* ≈ 1).

A simulation was run for each scenario consisting of 500 ASVs (*n* = 500), where the substrate feed concentration, *s*^*f*^, was set to 600 μM in an otherwise sterile medium (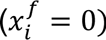), and the initial concentrations of *s*(0) and *x_i_* (0) were set to 0 and 10^−5^ μM, respectively. The dilution rate was stepped up from 0.1 to 1.0, and then 1.0 to 10 d^−1^ based on experimental event times (Table 1). We compare simulated bacterial ASVs, *x_i_*(*t*), partial pressure of oxygen in the gas headspace, *p*_*O*2_(*t*) (%), and dissolved oxygen, *O*_2_(*t*) (μM), to experimental results to determine if the specific growth rate distribution of prokaryotes in the chemostat is better described by uniform versus beta probability distributions. Simulations were conducted in Mathematica and the notebook and supporting materials are posted in a Github repository (https://github.com/maxEntropyProd/Trait-Based-Chemostat) and archived in Zenodo (doi: 10.5281/zenodo.10870761).

## Results

### Chemostat biogeochemistry

Chemostats were continuously monitored for carbon dioxide and oxygen in the headspace, and dissolved oxygen and pH in the culture. Percent CO_2_ averaged 0.09, 0.12, and 0.20%, percent O_2_ averaged 21.0, 20.9, and 20.5%, and dissolved O_2_ averaged 7.43, 7.10, and 5.18 mg L^−1^ for 0.1, 1, and 10 d^−1^ dilution rate periods, respectively (Fig. 2). Based on input and output O_2_ gas concentrations to each chemostat and steady state conditions, the rates of oxygen metabolism per liter of media were 0.171, 0.291, and 1.31 mmol O_2_ d^−1^ L^−1^ in MC1 and 0.175, 0.372, and 1.69 mmol O_2_ d^−1^ L^−1^ in MC2 for the 0.1, 1.0 and 10 d^−1^ dilution rate periods, respectively. Average pH was 7.29 for 0.1 d^−1^, 6.86 for 1 d^−1^, and 6.43 for 10 d^−1^. While community metabolism, based on DO, pH, O_2_, and CO_2_, was stable for most of the experiment, substantial instability occurred between days 21-24 in MC2 at a dilution rate of 10. d^−1^ as evident in the on-line measurements (Fig. 2, dashed lines).

**Fig. 2.**
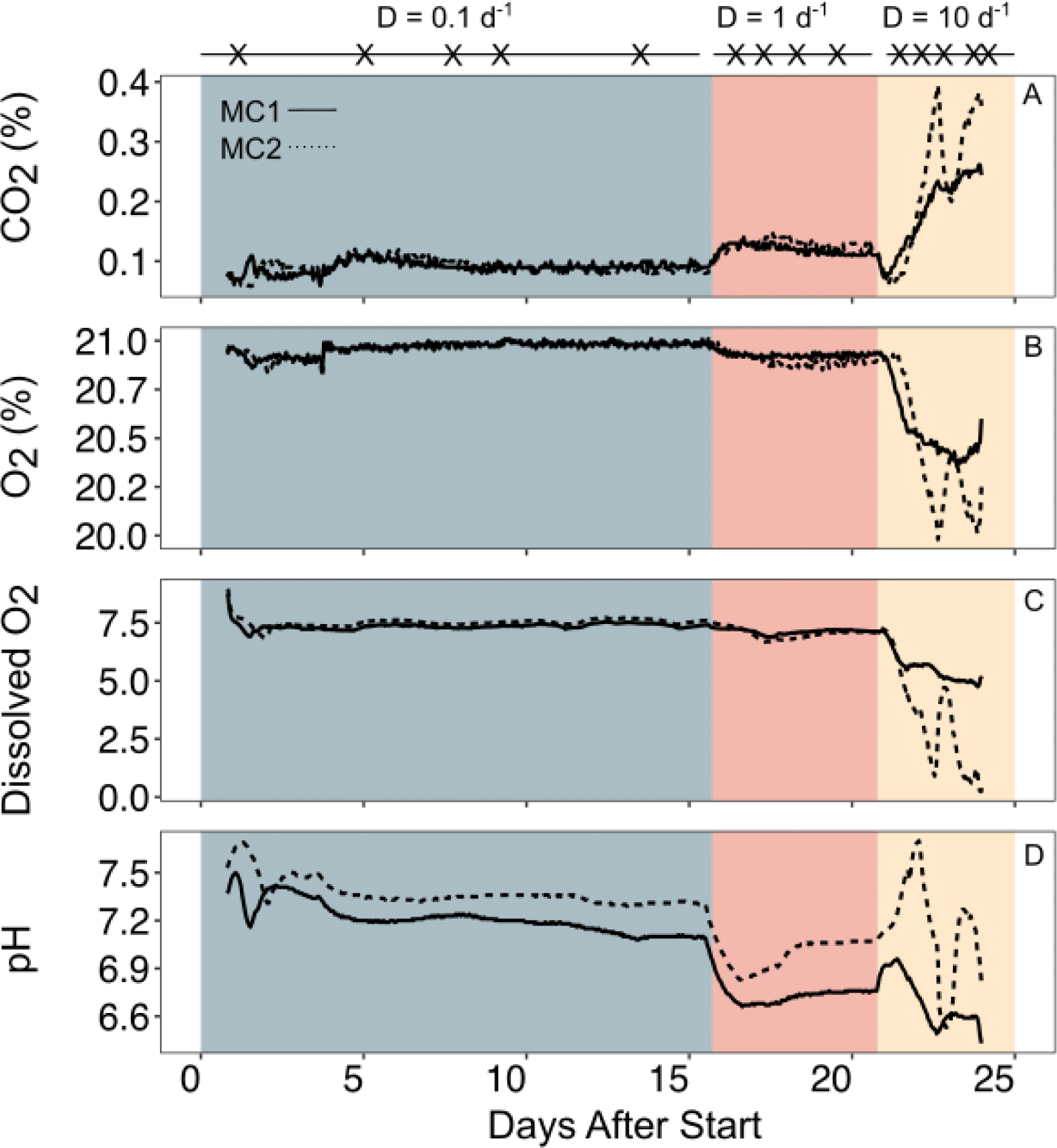
On-line measurements of (A) Carbon dioxide (%), (B) oxygen (%), (C) dissolved oxygen (mg L^−1^), and (D) pH for MC1 (solid line) and MC2 (dashed line), where background color indicates dilution rate. The sampling timeline (top) indicates when samples were collected for biogeochemical analyses and 16S rRNA gene sequencing.

Nitrate, ammonium, and phosphate started at initially high concentrations, reflecting the conditions in Siders Pond (Fig. 3). For both NO_3_^−^ and PO_4_^3−^, consumption rates (based on input-output concentrations and dilution rate) were significantly higher (p < 0.001) at 10 d^−1^ when compared to the 0.1 and 1.0 d^−1^ dilution periods. By the end of the low dilution rate period at 15.5 d, concentrations of all three inorganic nutrients had been reduced due to microbial growth and turnover of the initial pond water with feed media, and thus remained at low concentrations for the duration of the experiment. Dissolved organic carbon (DOC) concentration started low and remained so for the 0.1 d^−1^ dilution period, but then increased as dilution rate increased and the media CM:NM ratio (Ratio between the Carbon, CM, and Nutrient, NM, medium) decreased from 100:1 to 10:1 (Fig. 3D). DOC production was significantly higher in the 10 d^−1^ dilution period when compared to 0.1 d-^1^ (p = 0.01), but there was no significant difference between 1 d^−1^ and the other two periods.

**Fig. 3.**
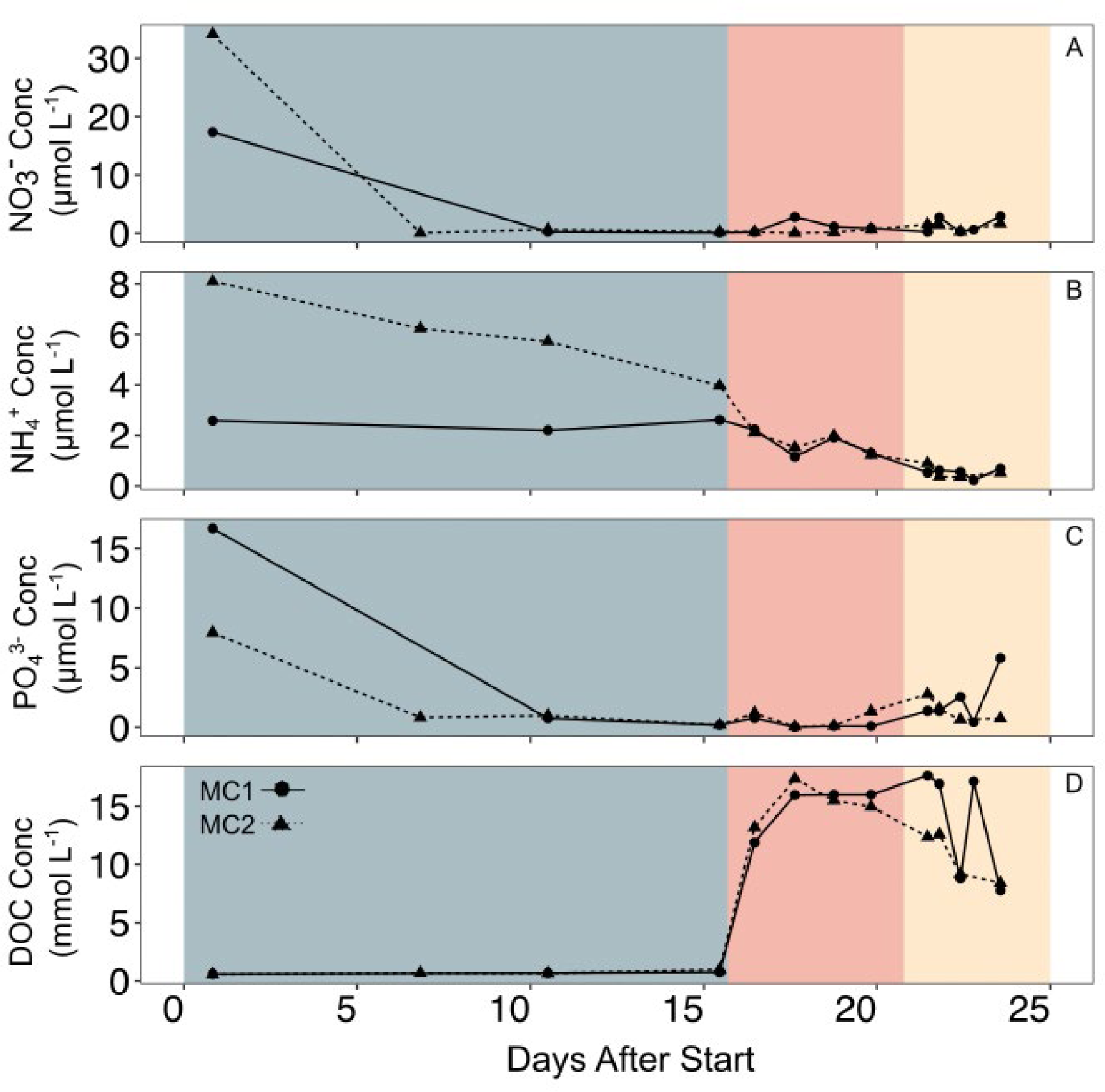
Concentrations (μM) of (A) nitrate, (B) ammonium, (C) phosphate, and (D) dissolved organic carbon (DOC) versus time. Solid lines with circles represent MC1, dashed lines with triangles represent MC2, and colors indicate 0.1 d^−1^ (blue), 1 d^−1^ (orange), and 10 d^−1^ (yellow) dilution rates, respectively.

### Chemostat microbial community

We sequenced 26 samples for the 16S rRNA gene representing 16 time points, including the initial pond sample, across the two MCs. After removing chimeras, 3,550,777 sequences remained, resulting in 1702 unique ASVs. After removing taxa matching Mitochondria or Chloroplasts, we were left with 1440 ASVs, 31 of which had a relative abundance greater than 5% in at least one sample (Fig. 4, Table S1). The sample collected from Sider’s Pond consisted of taxa belonging to families Saprospiraceae (∼20%), Rhodobacteraceae (∼16%), and Marinomonadaceae (∼6%), and was similar to the first sample of the 0.1 d^−1^ dilution treatment for both MC1 and MC2 at 0.8 d. This was except for Alteromonadaceae and Clade III (SAR11), both of which increased in relative abundance, and Saprospiraceae, which decreased in relative abundance, compared to the Siders sample. There were two samples from MC1 that did not successfully sequence (at 6.8 and 9.5 d) for the 0.1 d^−1^ dilution rate, but they did sequence in MC2 and were comprised primarily of Rhodobacteraceae (∼20-25%) and Methylophilaceae (genera *Methyloternera* ∼13-16% and *Methylophilus* ∼4-7%). Timepoint 10.5 d in MC1 looked like these two MC2 samples, except for an increase in Burkholderiaceae (∼8%). While MC1 at 15.5 d was like that of MC2 at 10.5 d, which was dominated by Flavobacteriaceae (∼51-55%), the two chemostats diverged in community composition from that point onward, even though external conditions where the same for both microcosms (Fig. 4).

**Fig. 4.**
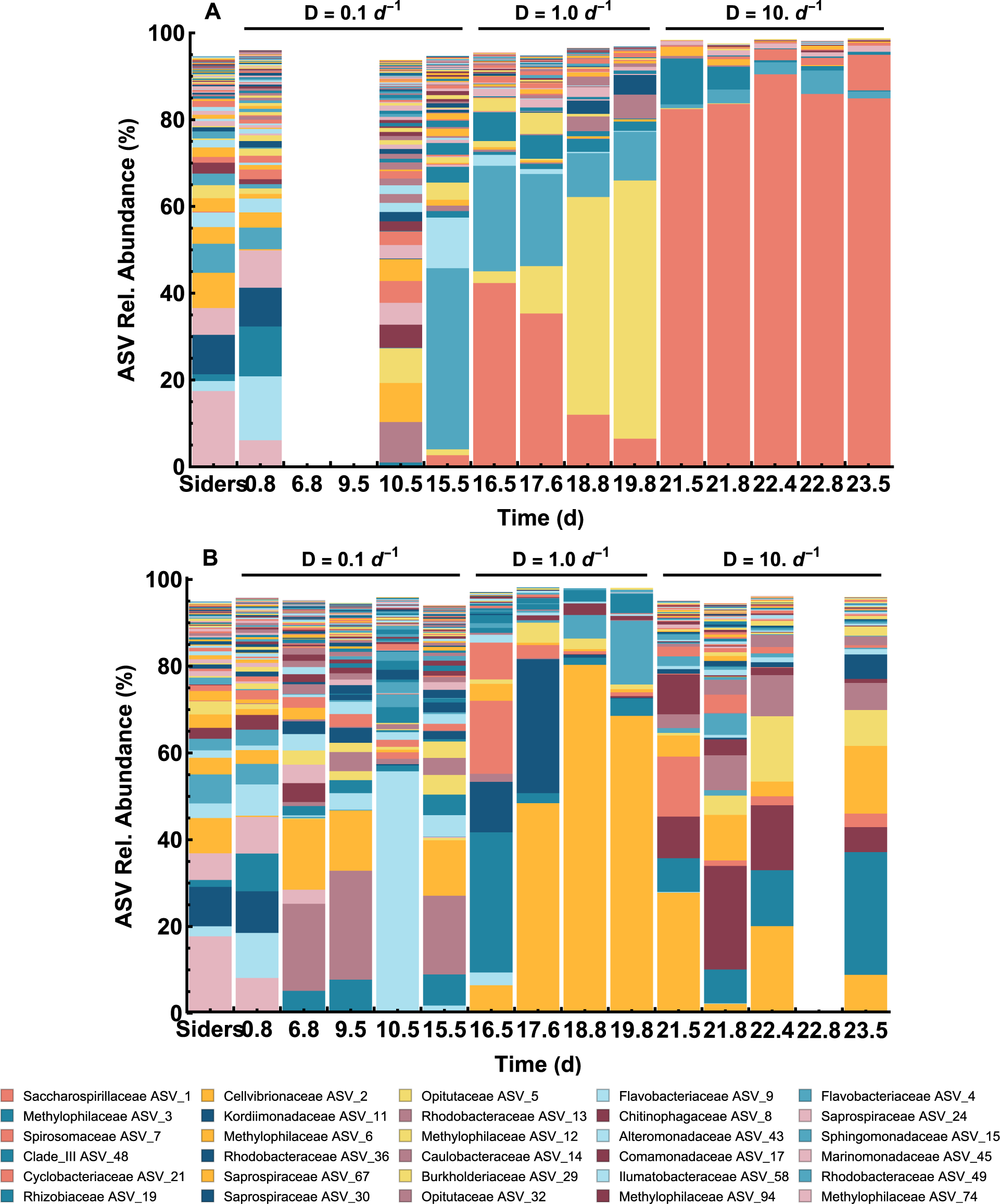
Relative abundance (%) of ASVs families from (A) MC1 and (B) MC2 versus time, along with ASVs from Siders Pond (“Siders” at time 0). Only ASVs that have an abundance greater than or equal to 0.1% are shown. The legend lists the top 30 ASVs families across all samples, and blank columns are for samples that did not sequence. The dilution rate and time of sample collection for each chemostat are indicated along the top and bottom, respectively.

There were clear successional patterns in MC1 throughout the 1 d^−1^ dilution rate. For the first two timepoints, Saccharospirillaceae (ASV 1; genus *Oceanobacter*), which made up ∼42% of the population on 16.5 d and ∼35% on 16.6 d, and Flavobacteriaceae, which made up ∼24% on 16.5 d and ∼21% on 17.6 d, were the most dominant families. Opitutaceae increased from ∼3% on 16.5 d up to ∼60% on 19.8 d, with Saccharospirillaceae and Flavobacteraceae decreasing to ∼6 and 11%, respectively. While both Rhizobiaceae and Paracaedicateraceae were present in 16.5 d and 17.6 d, they were effectively absent (<1%) by 18.6 d. We observed a similar successional pattern comprised of different ASVs in MC2 throughout the 1 d^−1^ dilution rate, where Methylophilaceae was the most dominant family (∼32%) on 16.5 d but was quickly displaced by Cellvibionaceae (∼48%) and Kordiimonadaceae (∼31%); Cellvibrionaceae then increased throughout the remainder of the 1 d^−1^ dilution rate sampling time points, making up ∼80% and 68% on 18.8 and 19.8 d, respectively (Fig. 4, Table S1).

Microbial community composition exhibited very different patterns once dilution rate increased to 10 d^−1^, and these patterns differed between chemostats. In MC1, the same ASV belonging to the Saccharospirallaceae family (ASV 1; genus *Oceanobacter*) increased from ∼82.5% on 21.5 d to 90.5% on 22.8 d, where it then decreased to 85% on 23.5 d due to an increase in a member of the Spirosomaceae family (ASV 7; ∼8%). The only other relatively dominant ASV belonged to the Methylophilaceae family (ASV 3), which also appeared for a short time in the lowest dilution rate (10.5 and 15.5 d) and re-appeared near the end of the 1 d^−1^ dilution rate (18.8 and 19.8 d). Cellvibrionaceae (ASV 2; ∼27.5%) continued to be the most dominant ASV in MC2 at the start of the 10 d^−1^ dilution rate (21.5 d), in addition to Spirosomaceae (ASV 7; ∼14%), Chitinophagaceae (ASV 8; ∼10%), Comamonadaceae (ASV 17; ∼9%), Methylophilaceae (ASVs 3 and 6; ∼8 and 5%), and Caulobacteraceae (ASV 14; ∼3%). Throughout the final three time points, Chitonophagaceae and Cellvibrionaceae demonstrated both increasing and decreasing patterns, while Methylophilaceae, composed of ASVs 3 (*Methylophilus*), ASV 6 (*Methylotenera*) and ASV 12 (*Methylophilus*), consistently increased to become the most dominant family (∼52%; Fig. 4; Table S1).

Mean cell abundance was 5.45 x 10^6^ ± 5.21 x 10^6^, 4.09 ± 10^6^ ± 1.88 x 10^6^, and 1.97 x 10^6^ ± 3.44 x 10^6^ cells L^−1^ for 0.1, 1, and 10 d^−1^ dilution rates, respectively (Fig. S1). There was a significant difference in beta dispersion among dilution rates in MC1 (Fig. 5A; *p* < 0.001, F_2,9_ = 17.93) but not in MC2 (Fig. 5B; *p* = 0.55). Average distance to group centroids decreased with increasing dilution rate for both chemostats, ranging from 0.53, 0.31, 0.14 (MC1) and 0.45, 0.34, 0.26 (MC2) for dilution rates 0.1, 1, and 10 d^−1^, respectively. Alpha diversity, represented here by Shannon indices, significantly decreased as dilution rate increased from 0.1 to 1.0 d^−1^ and from 1.0 to 10 d^−1^ for MC1 (Fig. 5C; p < 0.001, F_2,9_ = 6.76). Alpha diversity decreased between 0.1 to 1.0 d^−1^ for MC2, but not statistically between 1.0 and 10. d^−1^, where it increased (p = 0.002, F_2,10_ = 12.95). Community stability (i.e., community similarity between consecutive time points) increased significantly between the 0.1 and 1.0 d^−1^ dilution rate for MC1, but not between 1.0 and 10 d^−1^ (Fig. 5D; p = 0.002, F_2,6_ = 21.54), and there was no statistically significant trend for MC2 (p = 0.14). Differences among groups were significant (*p* < 0.001, F_2,16_ = 13.36) between 0.1 d^−1^ and 1 d^−1^ dilution rates, but not between 1 d^−1^ and 10 d^−1^.

**Fig. 5.**
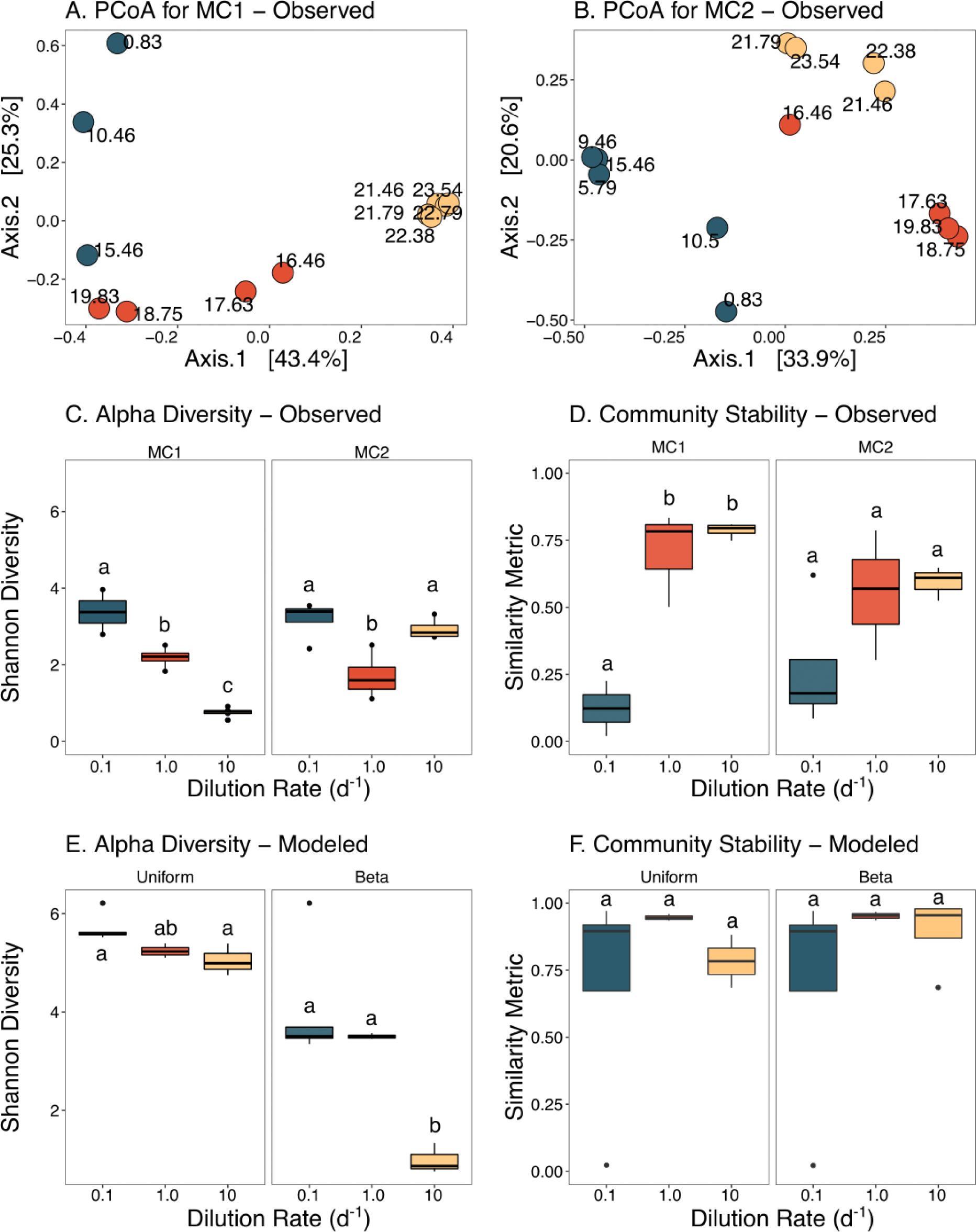
Ordination of principal coordinates analysis based on Bray-Curtis similarity among samples for (A) MC1 and (B) MC2, where numbers represent time (in days) after the start of the experiment. Boxplots represent (C) alpha diversity based on Shannon Index and (D) community stability based on Bray-Curtis similarity between consecutive time points separated by MC, as well as (E) alpha diversity and (F) community stability for modeled data from the uniform (left) beta (right) distribution simulation. Groups that do not share the same letter are significantly different according to a Tukey’s HSD multiple comparison test. Color indicates dilution rate (blue, 0.1 d^−1^; red, 1.0 d^−1^; yellow, 10. D^−1^).

### Simulations

In the simulation run where the growth efficiency traits (*ε*_*i*_) were drawn from a uniform probability distribution (Fig. 1A), ASV diversity was high at all time points that correspond to MC sample times (Fig. 6A), and alpha diversity averaged over each dilution rate (Fig. 5E) was higher than observed for all dilutions rates (Fig. 5C). There was only a small decrease in diversity as dilution rate increased. In the second scenario, where *ε*_*i*_ was drawn from a beta probability distribution (Fig. 1B), diversity was greatly reduced for all three dilution rates, and there was a significant loss of diversity at the highest dilution rate (Fig. 5E; p = 0.04, *χ*^2^ = 6.27, df = 2), which is also evident in comparing the two timeseries plots (Fig. S2). Both simulations show that populations are more stable at each dilution rate as compared to the observed data, especially at D = 0.1 d^−1^ (Figs. 5F vs 5D). The total bacterial biomass between the two simulations were similar (Fig. S3), but the model shows increasing biomass as dilution rate increases, while the trend is the opposite for the observed cell counts (Fig. S1). In general, there is close agreement between modeled and observed oxygen concentrations in both the headspace (pO2) and medium (DO) for both simulations (Fig. S4); however, the significant fluctuations in headspace O_2_ and media DO in MC2 (Fig. S4, gray lines) at the highest dilution rate was not captured by either simulation, which indicates some instability in community metabolism in MC2 at D = 10. d^−1^. This instability is also evident in the CO_2_ observations (Fig. 2A, dashed line).

**Fig. 6.**
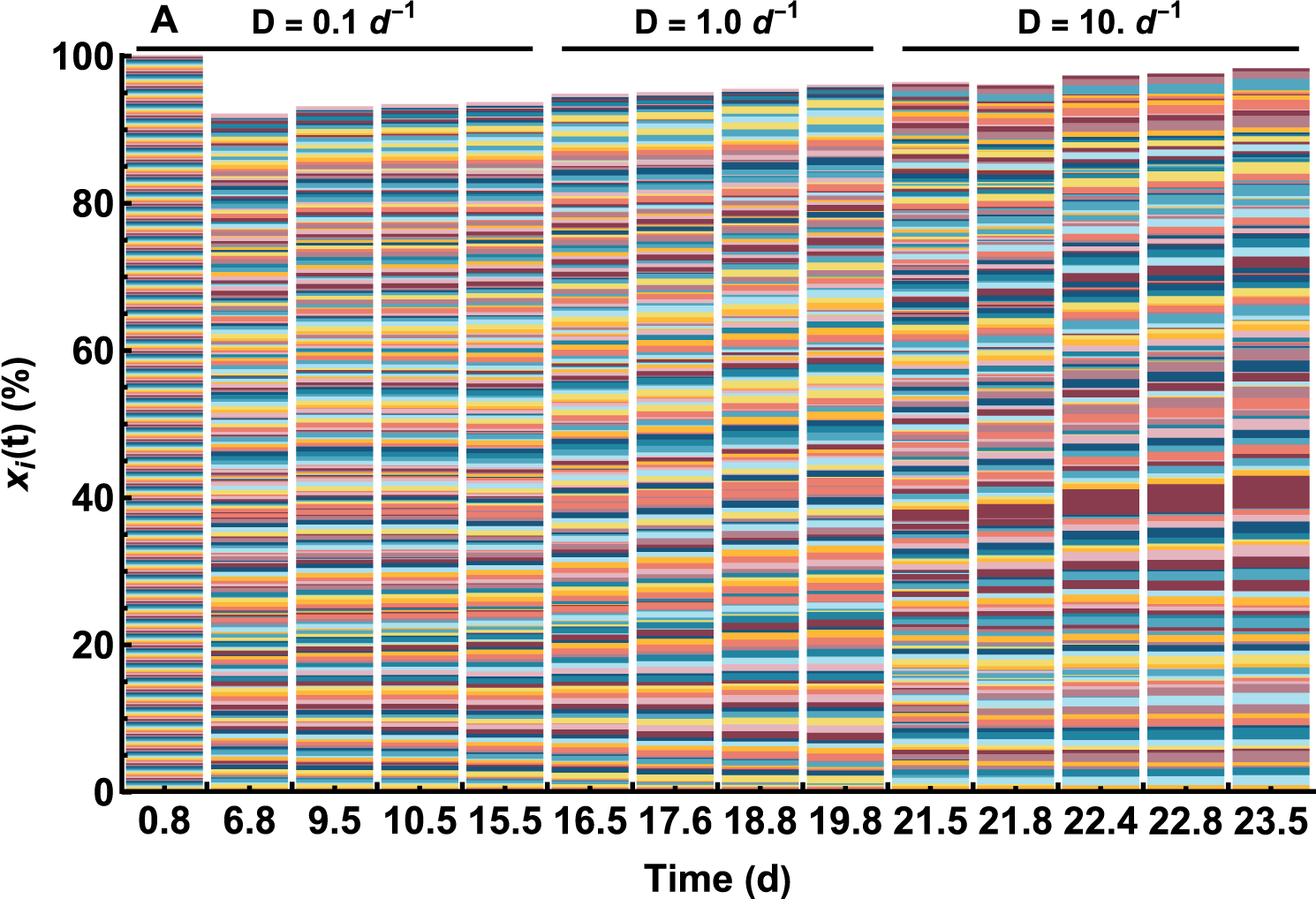
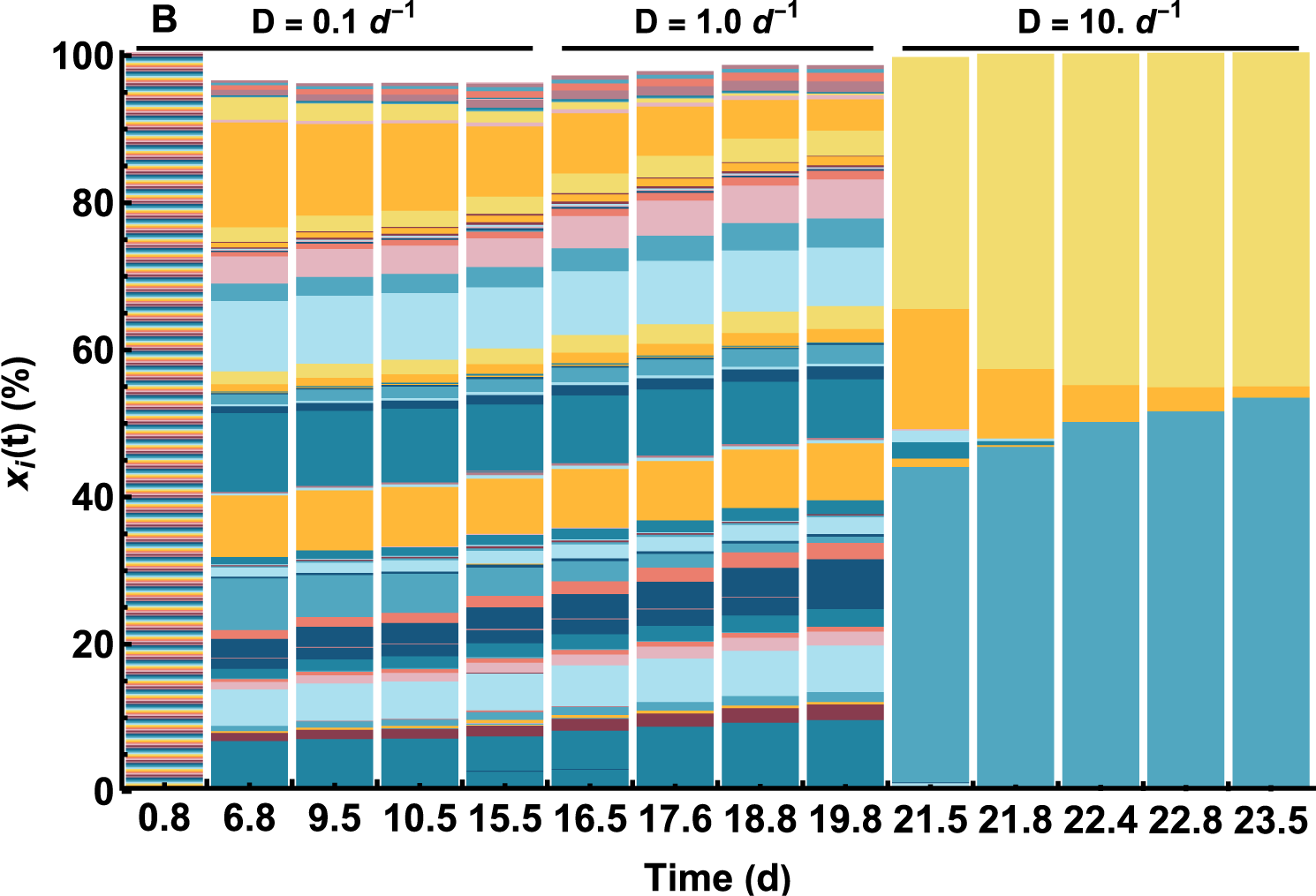
Relative abundance (%) of simulated ASVs at selected times for *εi* drawn from (A) uniform and (B) beta probability distributions (see Fig. 1). Only ASVs whose relative abundance equals or exceeds 0.1% are shown. Dilution rate at time of model query is shown at the top.

## Discussion

Here we discuss how dilution rate (or water turnover rate in the chemostats) affects microbial community composition, the importance of endogenous versus exogenous drivers, and, briefly, connections between biogeochemistry and community stability, or the lack thereof.

When operating at the lowest dilution rate (0.1 d^−1^), alpha diversity was highest compared to higher dilution rates (Fig. 5C). Low dilution rate, where resources were turned over only once every 10 days and concentrations were low (Fig. 3, t > 10 d), favors bacteria with gleaner-type behavior characterized by high affinity for resources but low growth rate. These experimental conditions supported a high diversity of bacteria, but it is unclear if the high diversity is due to co-existence mechanisms that prevented competitive exclusion from occurring, or if bacterial washout was not given sufficient time to occur; however, we have run similar chemostat experiments at 0.1 d^−1^ dilution rate that maintained high diversity for many years, so the latter seems unlikely (Fernandez-Gonzalez et al. 2016). Simulation results at the lowest dilution rate between the uniform and beta distributions of *ε*_*i*_ both show high diversity (Fig. 5E and Fig. 6, times 0.8 to 15.5 d); however, diversity using the beta distribution model for *εi* is closer to that observed for both chemostats (Fig. 5C vs 5E).

When dilution rate was increased to 1 d^−1^, alpha diversity decreased in both chemostats and a few ASVs became more dominant over time - Optitutaceae (ASV 5) and Cellvibrionaceae (ASV 2) in MC1 and MC2, respectively (Fig. 4). This suggests that these taxonomic groups exist intermediately along the gleaner-opportunist continuum, with a slight competitive advantage of these taxa resulting in higher abundances at the expense of others. Our model results, however, still show relatively high diversity for both the uniform and beta initial distributions for *ε*_*i*_, but the diversity is lower in the simulation based on the beta distribution compared to the uniform one (Fig. 5E and Fig. 6, times 16.5 to 19.8 d).

At a dilution rate of 10 d^−1^, *Oceanobacter* (ASV 1) dominated MC1 (>80% of the community), suggesting this taxonomic group is capable of fast enough growth to avoid being washed out of the chemostat. However, MC2 exhibited different behavior in that diversity increased from that observed at D = 1.0 d^−1^ (Fig. 5C). It appears that MC2 experienced some kind of metabolic instability, as large oscillations in pO_2_, DO, CO_2_ and pH were evident during this phase (Fig. 2, t > 21 d, dashed lines), which may explain the higher diversity in MC2 at 10 d^−1^ rate because the chemostat was not near steady-state operation. The genera *Methylophilus* (ASV 3 and 12) and *Methylotenera* (ASV 6) of the Methylophilaceae family comprised more than half of the community during the transient in MC2 (Fig. 4B) and are known methylotrophs (Doronina et al. 2014), so perhaps there was a transient shift to methylotrophy in MC2. Although most methylotrophs are slow growers (<1 d^−1^), there are some with specific growth rates greater than 10 d^−1^ (Silman et al. 1989). Because of the functional instability in MC2 at D = 10 d^−1^, we will remove it from our discussion below because it did not reach functional steady state.

### Diversity decreases at the growth rate boundary

In our experiment and model based on the beta probability distribution of *εi* that is skewed towards oligotrophic growth, alpha diversity significantly decreased as dilution rate increased from 0.1 d^−1^ to 10 d^−1^ (Figs. 5C and 5E). We argue this result is rooted in the ‘boundary of life’ concept, where species diversity decreases as the boundaries where life can exist are approached – in this case, maximum specific growth rate. When we draw *εi* from a uniform distribution, community diversity remained high even at the 10 d^−1^ dilution rate, which is inconsistent with our observations. To persist in the chemostat, a bacterium’s net growth rate must equal or exceed the dilution rate to avoid being washed out of the system. As dilution rate increases, only those bacteria with high enough net specific growth rates will persist. Results show that only a small proportion of the natural pond community could grow fast enough to persist under the highest dilution rate imposed, which is consistent with Weisssm et al. (2010), who found a bias for extremely slow-growing organisms based on codon usage patterns from genomic surveys. In a resource competition context, the functional traits most relevant are maximum specific growth rate and substrate affinity, as these two strategies are among the most important mechanistic drivers of biodiversity (Chesson 2000). However, there is a trade-off between substrate affinity (or low minimum resource requirement) and maximum specific growth rate (Grover 1997, Follows et al. 2007). Gleaners (or K-strategists) invest in metabolic capability that allows for high substrate affinity at the expense of high specific growth rates, but this gives them an advantage when substrate concentrations are low. Opportunists (or r-strategists) allocate metabolic capability towards high specific growth rate but sacrifice substate affinity, which allows them to outcompete gleaners, but only when substrate concentrations are high (Norris et al. 2020). This trade-off is not binary but is a continuum between the two extremes, so it is represented with the trait variable *ε*_*i*_ in our adaptive Monod kinetics model (Eq. 1) that varies between 0 (gleaner) and 1 (opportunist). These differing growth strategies likely shaped the changes in observed microbial diversity as a function of dilution rate, where resource competition changed from a high diversity gleaner-type community at the 0.1 d^−1^ dilution rate to a low diversity opportunist-like community at the 10 d^−1^ dilution rate, since resource concentrations increase with increasing dilution rate (Smith & Waltman 1995).

### Increased influence of exogenous driver dampens community dynamics

Several studies have demonstrated inherently unstable microbial community compositions over space and time, with examples including plankton (Benincà et al. 2009, Needham & Fuhrman 2016), methanogenic (Fernandez et al. 2000), methanotrophic (Fernandez-Gonzalez et al. 2016), and nitrifying (Graham et al. 2007) communities. Similarly, we also observed changes in dynamics and patterns of community turnover as a function of dilution rate as well as changes in diversity. In both chemostats, community stability increased (Fig. 5D) and distance-to-group centroid decreased between sampling time points with increasing dilution rate (Figs. 5A, B). We attribute this pattern to a shift from endogenous dynamics as the primary driver of community composition (Konopka et al. 2015), to exogenous forcing as dilution rate increased from 0.1 to 10 d^−1^. For example, if the dilution rate was dropped to 0 d^−1^, then external forcing would be effectively removed so that community population dynamics would be a result of entirely endogenous dynamics. As external flow is imposed and dilution rate increases, competitive exclusion (Hardin 1960) often does not occur as community diversity remains high for extended periods (Fernandez-Gonzalez et al. 2016), and the so called “paradox of the plankton” (Hutchinson 1961) can be observed in the laboratory or natural systems. There are numerous theories that have been proposed to explain why diversity remains high, which are often associated with rapid turnover of a community when external forcing is weak (Chesson 2000). Some of these theories include kill-the-winner, where viruses lyse the dominate taxa (Thingstad & Lignell 1997), cross-feeding that can maintain diversity as well as lead to instabilities (Liu & Sumpter 2017, Butler & O’Dwyer 2018), and niche creation (Callahan et al. 2014), but there are many others (Louca et al. 2018).

As dilution rate is increased further, however, system heterogeneity can be harder to maintain, and metabolites associated with cross-feeding can be diluted or effectively washed out of the system. Furthermore, competitive exclusion becomes more relevant on the experimental time scale of study. At the extreme, when the dilution rate crosses the boundary defined by the maximum specific growth rate of the community, all organisms are washed out of the chemostat, and the system becomes the same as the input feed. At this point the system matches the exogenous driver. If communities are skewed towards oligotrophic growth characteristics, as we propose here, then near the growth rate boundary, species diversity becomes depleted, making the system even more stable as fewer organisms are present for succession, which might be occurring in the chemostats at the 10 d^−1^ dilution rate. High system turnover becomes particularly relevant for structuring communities.

Recently, Ratzke et al. (2020) examined how resource concentration alters community diversity and stability, but under conditions closer to batch operating conditions that lack exogenous forcing. Under high initial nutrient concentration, they found that the community lost diversity, while low initial nutrient concentrations resulted in communities with high diversity. They concluded that high nutrient availability leads to high bacterial uptake, which subsequently alters the local environment, such as changing pH or dissolved oxygen concentration, which leads to death and loss of diversity as those boundaries are approached. These results are similar to ours, in that bacterial growth in eutrophic environments drives the system towards the habitable limit, such as extreme pH or low dissolved oxygen that fewer species are capable of growing at. This results in loss of diversity in a manner similar to high dilution rates. Ratzke et al. also conducted chemostat-like experiments (periodically diluted batch growth experiments), but instead of altering the dilution rate, they changed the nutrient concentrations in the feed between either low or high concentrations. They found low nutrient concentration produced stable communities, but high nutrient concentration resulted in unstable communities driven by fluctuations in pH. These results differ from ours, in that we found unstable dynamics at low dilution rate (low nutrient concentration and supply rate), but stable communities at high dilution rate (high nutrient concentration and supply rate). However, the two experiments differ in that exogenous forcing by high dilution rate may provide stability that is absent in their periodically fed batch experiments. Further studies are needed to properly assess the differences between these two approaches.

As noted by Locey and Lennon (2019), there have been relatively few studies relating residence time (τ; the inverse of dilution rate, *DD*) to biodiversity and related metrics of community assembly. Locey and Lennon (2019) used a trait-based model study that drew from a uniform distribution and found that species richness decreased with increasing dilution rate as we did, but they also found at dilution rates lower than ∼10^−3^ d^−1^, richness began to decrease, thereby creating a unimodal distribution in richness as a function of *DD*. Mansfeldt et al. (2019) investigated the impact of microbial residence time (MRT) on community composition (16S rRNA-based) and function (RNAseq-based) in laboratory sequencing batch reactors treating wastewater over residence times varying from 1 to 15 days (or dilution rates from 0.066 to 1 d^−1^). Similar to our study, both the experimental results and the model they developed showed a decrease in Shannon diversity and richness as dilution rate increased. Both the Locey and Lennon (2019) and Mansfeldt et al. (2019) studies differ from ours in objectives and design making them difficult to compare, but all suggest that traits, such as the maximum specific growth rate, are not uniformly distributed in a community but are instead skewed.

### Biogeochemical consequences

Finally, we note that although the communities exhibited considerable turnover of ASVs, particularly at low and intermediate dilution rates, these dynamics are not particularly evident in the biogeochemical processes, at least as observed in respiration (CO_2_ production and O_2_ consumption, Fig. 2, S4), pH, and nutrient concentrations (Fig. 3), except in MC2 during the highest dilution rate. Functional stability despite unstable community composition has been noted by others (Fernandez et al. 1999, Fernandez-Gonzalez et al. 2016), and is the basis for ITSNTS theory (“It’s the song not the singer”). Doolittle and Inkpen (2018) propose that biogeochemical processes are the unit of selection, where biogeochemistry is the “song” and a particular subset of the microbial community are the “singers”, but multiple configurations can maintain the song through functional equivalencies (Louca et al. 2018). Under such a perspective, stability of the community is not required, nor should community instability necessarily be a point of concern. However, as the growth rate boundary is approached and the diversity of the community decreases, the ability of the system to maintain biogeochemical function eventually becomes compromised, although the community, or individual, may be stable. In ITSNTS parlance, the song’s fidelity may be compromised with insufficient singers. Perhaps this is the cause of the functional instability in MC2 at D = 10 d^−1^.

## Conclusion

We conducted a chemostat experiment using a natural pond microbial community in conjunction with an adaptive Monod trait-based model to examine how community diversity changes as a function of specific growth rate dictated by the chemostat dilution (or turnover) rate. We hypothesized that the number of bacterial species (alpha diversity) would decrease under increasing dilution rates, because natural microbial communities are skewed towards bacteria with oligotrophic rather than copiotrophic growth characteristics. A result of this skewedness is that the dynamics of the community decreased with increasing dilution rate because of the loss of functional redundancy. The rapid community turnover that we have observed at lower dilution rate in this study, and in other chemostat experiments (Fernandez-Gonzalez et al. 2016), decreased due to the lack of diversity at high dilution rates. Model results show that when the trait variable is drawn from a beta distribution that is skewed towards oligotrophic growth strategies, alpha diversity decreases and community stability increases with increasing dilution rate as observed in the chemostats, supporting the skewed growth rates hypothesis. When the trait variable is drawn from a uniform distribution, communities retain high diversity as the upper growth rate boundary is approached, which is inconsistent with our observations. These results also illustrate the importance of trait-based model initialization for proper prediction of biodiversity dynamics in microbial communities.

## Acknowledgments

This research was supported by grants from NSF #1655552 (JJV and JAH), #1637630 (JJV), and #1907285 (ANB), and by the Simons Foundation CBIOMES project #549941FY22 (JJV). Undergraduate Yang Wenzhou was supported by MBL’s Semester in Environmental Science Program.

## Supplemental Figures

**Fig. S1.**
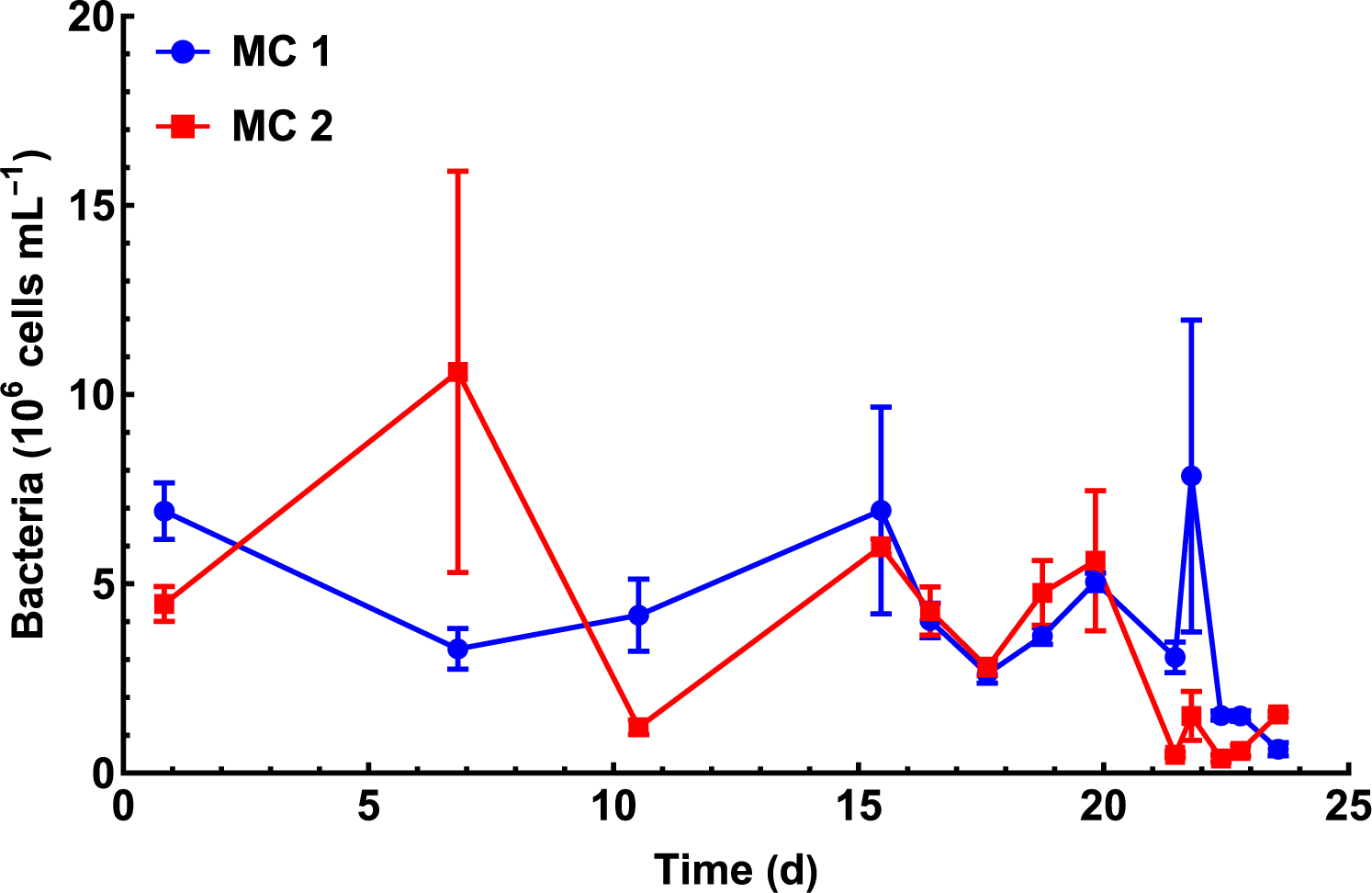
Bacterial cell counts form MC 1 (blue line) and MC 2 (red line) over time. Error bars represent standard error.

**Fig. S2.**
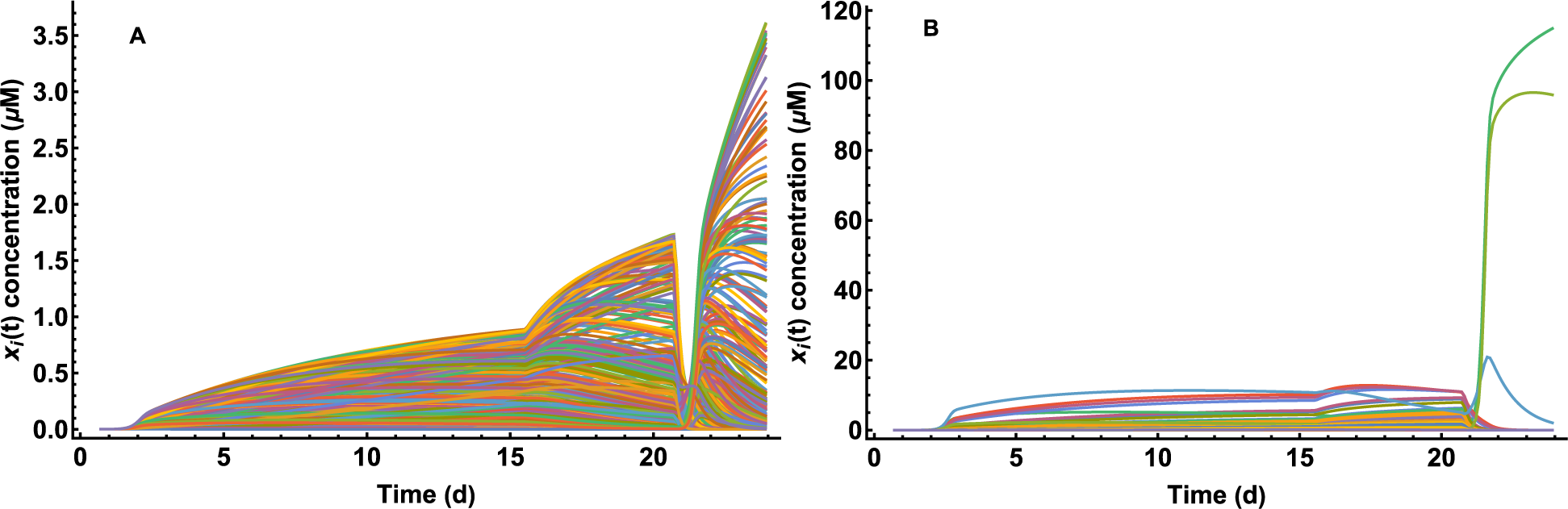
Simulated ASV absolute concentrations (μM C) versus time for *εi* drawn from (A) uniform and (B) beta probability distributions (See Fig. 1).

**Fig. S3.**
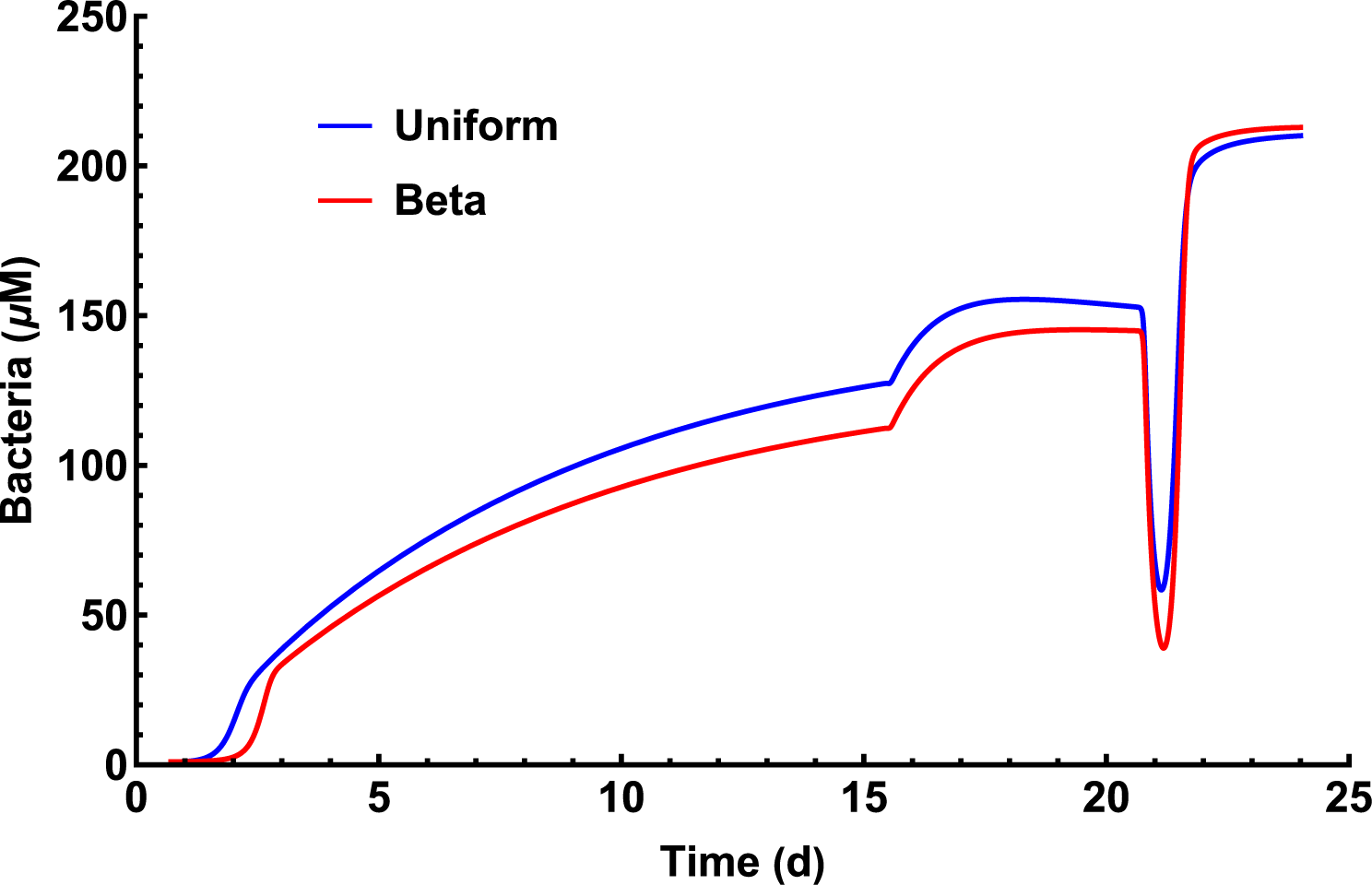
Simulated bacterial concentration (μM C) versus time (d) for *εi* drawn from uniform (blue line) and beta (red line) probability distributions.

**Fig. S4.**
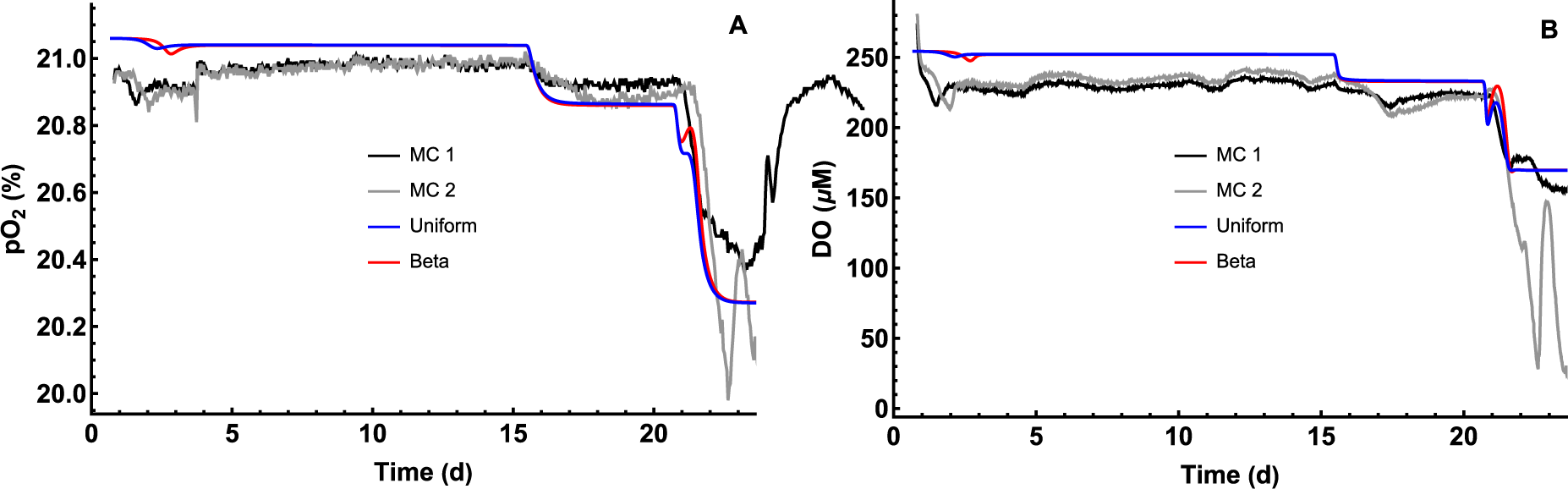
Simulated (blue and red lines) and observed (black and gray lines) (A) partial pressure of oxygen in the chemostat headspace (pO_2_, %) and (B) dissolved oxygen concentration in the media (DO, μM) versus time (d).

## References

Alberty, R. A. 2003. Thermodynamics of biochemical reactions. Wiley & Sons, Hoboken, NJ.

Algar, C. K., and J. J. Vallino. 2014. Predicting microbial nitrate reduction pathways in coastal sediments. Aquat.Microb.Ecol. 71:223–238, doi:10.3354/ame01678.

Baker, B. J., and J. F. Banfield. 2003. Microbial communities in acid mine drainage.FEMS Microbiology Ecology 44:139–152, doi:10.1016/s0168-6496(03)00028-x.

Benincà, E., K. D. Jöhnk, R. Heerkloss, and J. Huisman. 2009. Coupled predator–prey oscillations in a chaotic food web. Ecology Letters 12:1367–1378, doi:10.1111/j.1461-0248.2009.01391.x.

Bhaya, D. 2012. Probing Functional Diversity of Thermophilic Cyanobacteria in Microbial Mats. Pages 17–46 in R. Burnap and W. Vermaas, editors. Functional Genomics and Evolution of Photosynthetic Systems. Springer Netherlands, Dordrecht, doi:10.1007/978-94-007-1533-2_2.

Butler, S., and J. P. O’Dwyer. 2018. Stability criteria for complex microbial communities. Nature Communications 9:10, doi:10.1038/s41467-018-05308-z.

Callahan, B. J., T. Fukami, and D. S. Fisher. 2014. Rapid evolution of adaptive niche construction in experimental microbial populations. Evolution 68:3307–3316, doi:10.1111/evo.12512.

Caraco, N. 1986. Phosphorus, iron, and carbon cycling in a salt stratified coastal pond. Boston University, Boston.

Caraco, N., A. Tamse, O. Boutros, and I. Valiela. 1987. Nutrient limitation of phytoplankton growth in brackish coastal ponds. Canadian Journal of Fisheries and Aquatic Sciences 44:473–476, doi:10.1139/f87-056.

Chesson, P. 2000. Mechanisms of maintenance of species diversity. Annual Review of Ecology and Systematics 31:343–366, doi:10.1146/annurev.ecolsys.31.1.343.

Doolittle, W. F., and S. A. Inkpen. 2018. Processes and patterns of interaction as units of selection: An introduction to ITSNTS thinking. Proceedings of the National Academy of Sciences 115:4006–4014, doi:10.1073/pnas.1722232115.

Doronina, N., E. Kaparullina, and Y. Trotsenko. 2014. The Family Methylophilaceae. Pages 869–880 in E. Rosenberg, E. F. DeLong, S. Lory, E. Stackebrandt, and F. Thompson, editors. The Prokaryotes: Alphaproteobacteria and Betaproteobacteria. Springer Berlin Heidelberg, Berlin, Heidelberg, doi:10.1007/978-3-642-30197-1_243.

Falkowski, P. G., T. Fenchel, and E. F. DeLong. 2008. The Microbial Engines That Drive Earth’s Biogeochemical Cycles. Science 320:1034–1039, doi:10.1126/science.1153213.

Fernandez-Gonzalez, N., J. A. Huber, and J. J. Vallino. 2016. Microbial Communities Are Well Adapted to Disturbances in Energy Input. mSystems 1:1–15, doi:10.1128/mSystems.00117-16.

Fernandez, A., S. Huang, S. Seston, J. Xing, R. Hickey, C. Criddle, and J. Tiedje. 1999. How Stable Is Stable? Function versus Community Composition. Applied and Environmental Microbiology 65:3697–3704.

Fernandez, A. S., S. A. Hashsham, S. L. Dollhopf, L. Raskin, O. Glagoleva, F. B. Dazzo, R. F. Hickey, C. S. Criddle, and J. M. Tiedje. 2000. Flexible community structure correlates with stable community function in methanogenic bioreactor communities perturbed by glucose. Appl.Microbiol.Biotechnol. 66:4058–4067.

Follows, M. J., and S. Dutkiewicz. 2011. Modeling Diverse Communities of Marine Microbes. Annual Review of Marine Science 3:427–451, doi:10.1146/annurev-marine-120709-142848.

Follows, M. J., S. Dutkiewicz, S. Grant, and S. W. Chisholm. 2007. Emergent Biogeography of Microbial Communities in a Model Ocean. Science 315:1843–1846, doi:10.1126/science.1138544.

Frank, K. L. 2013. Linking metabolic rates with the diversity and functional capacity of endolithic microbial communities within hydrothermal vent structures. Harvard University.

Fredrickson, A., and G. Stephanopoulos. 1981. Microbial competition. Science 213:972–979, doi:10.1126/science.7268409.

Graham, D. W., C. W. Knapp, E. S. Van Vleck, K. Bloor, T. B. Lane, and C. E. Graham. 2007. Experimental demonstration of chaotic instability in biological nitrification. ISME J 1:385–393, doi:10.1038/ismej.2007.45.

Green, J. L., B. J. M. Bohannan, and R. J. Whitaker. 2008. Microbial Biogeography: From Taxonomy to Traits. Science 320:1039–1043.

Grover, J. P. 1990. Resource Competition in a Variable Environment: Phytoplankton Growing According to Monod’s Model. The American Naturalist 136:771–789, doi:10.1086/285131.

Grover, J. P. 1997. Resource competition. Springer Science & Business Media.

Hardin, G. 1960. The Competitive Exclusion Principle. Science 130:1292–1297.

Holling, C. S. 1965. The functional response of predators to prey density and its role in mimicry and population regulation. Mem.Entom.Soc.Can. 45:1–60.

Hutchinson, G. E. 1961. The paradox of the plankton. Amer.Nat. 95:137–145.

Inskeep, W., Z. Jay, S. Tringe, M. Herrgard, and D. Rusch. 2013. The YNP Metagenome Project: Environmental Parameters Responsible for Microbial Distribution in the Yellowstone Geothermal Ecosystem. Frontiers in Microbiology 4, doi:10.3389/fmicb.2013.00067.

Konopka, A., S. Lindemann, and J. Fredrickson. 2015. Dynamics in microbial communities: unraveling mechanisms to identify principles. ISME J 9:1488–1495, doi:10.1038/ismej.2014.251.

Krause, S., X. Le Roux, P. A. Niklaus, P. M. Van Bodegom, J. T. Lennon, S. Bertilsson, H.-P. Grossart, L. Philippot, and P. L. E. Bodelier. 2014. Trait-based approaches for understanding microbial biodiversity and ecosystem functioning. Frontiers in Microbiology 5, doi:10.3389/fmicb.2014.00251.

Lajoie, G., and S. W. Kembel. 2019. Making the Most of Trait-Based Approaches for Microbial Ecology. Trends in Microbiology 27:814–823, doi:10.1016/j.tim.2019.06.003.

Lewin, A., A. Wentzel, and S. Valla. 2013. Metagenomics of microbial life in extreme temperature environments. Current Opinion in Biotechnology 24:516–525, doi:10.1016/j.copbio.2012.10.012.

Li, J., S. Browning, S. P. Mahal, A. M. Oelschlegel, and C. Weissmann. 2010. Darwinian Evolution of Prions in Cell Culture. Science 327:869–872, doi:10.1126/science.1183218.

Litchman, E., K. F. Edwards, and C. A. Klausmeier. 2015. Microbial resource utilization traits and trade-offs: implications for community structure, functioning, and biogeochemical impacts at present and in the future. Frontiers in Microbiology 6:1–10, doi:10.3389/fmicb.2015.00254.

Liu, Y., and D. Sumpter. 2017. Insights into resource consumption, cross-feeding, system collapse, stability and biodiversity from an artificial ecosystem. Journal of The Royal Society Interface 14:1–12, doi:10.1098/rsif.2016.0816.

Locey, K. J., and J. T. Lennon. 2019. A Residence Time Theory for Biodiversity. The American Naturalist 194:59–72, doi:10.1086/703456.

Louca, S., M. F. Polz, F. Mazel, M. B. N. Albright, J. A. Huber, M. I. O’Connor, M. Ackermann, A. S. Hahn, D. S. Srivastava, S. A. Crowe, M. Doebeli, and L. W. Parfrey. 2018. Function and functional redundancy in microbial systems. Nature Ecology & Evolution, doi:10.1038/s41559-018-0519-1.

Lynch, M. D. J., and J. D. Neufeld. 2015. Ecology and exploration of the rare biosphere. Nat Rev Micro 13:217–229, doi:10.1038/nrmicro3400.

Macarthur, R. H., and E. O. Wilson. 1967. The Theory of Island Biogeography. REV - Revised edition. Princeton University Press.

Malik, A. A., J. B. H. Martiny, E. L. Brodie, A. C. Martiny, K. K. Treseder, and S. D. Allison. 2020. Defining trait-based microbial strategies with consequences for soil carbon cycling under climate change. The ISME Journal 14:1–9, doi:10.1038/s41396-019-0510-0.

Mansfeldt, C., S. Achermann, Y. Men, J.-C. Walser, K. Villez, A. Joss, D. R. Johnson, and K. Fenner. 2019. Microbial residence time is a controlling parameter of the taxonomic composition and functional profile of microbial communities. The ISME Journal 13:1589–1601, doi:10.1038/s41396-019-0371-6.

McMurdie, P. J., and S. Holmes. 2013. phyloseq: An R Package for Reproducible Interactive Analysis and Graphics of Microbiome Census Data. PLoS ONE 8:1–11, doi:10.1371/journal.pone.0061217.

Needham, D. M., and J. A. Fuhrman. 2016. Pronounced daily succession of phytoplankton, archaea and bacteria following a spring bloom. Nature Microbiology 1:1–7, doi:10.1038/nmicrobiol.2016.5.

Norris, N., N. M. Levine, V. I. Fernandez, and R. Stocker. 2020. Mechanistic model of nutrient uptake explains dichotomy between marine oligotrophic and copiotrophic bacteria. bioRxiv:2020.2010.2008.331785, doi:10.1101/2020.10.08.331785.

Oksanen, J., F. G. Blanchet, M. Friendly, R. Kindt, P. Legendre, D. McGlinn, P. Minchin, R. O’Hara, G. Simpson, P. Solymos, M. Stevens, E. Szoecs, and H. Wagner. 2019. Vegan: Community ecology package https://cran.r-project.org/web/packages/vegan/index.html.

Porter, K. G., and Y. S. Feig. 1980. The use of DAPI for identifying and counting aquatic microflora. Limnol.Oceanogr. 25:943–948.

Ratzke, C., J. Barrere, and J. Gore. 2020. Strength of species interactions determines biodiversity and stability in microbial communities. Nature Ecology & Evolution 4:376–383, doi:10.1038/s41559-020-1099-4.

Rothschild, L. J., and R. L. Mancinelli. 2001. Life in extreme environments. Nature 409:1092–1101.

Rousk, J., E. Bååth, P. C. Brookes, C. L. Lauber, C. Lozupone, J. G. Caporaso, R. Knight, and N. Fierer. 2010. Soil bacterial and fungal communities across a pH gradient in an arable soil. The ISME Journal 4:1340–1351, doi:10.1038/ismej.2010.58.

Sharp, C. E., A. L. Brady, G. H. Sharp, S. E. Grasby, M. B. Stott, and P. F. Dunfield. 2014. Humboldt’s spa: microbial diversity is controlled by temperature in geothermal environments. The ISME Journal 8:1166–1174, doi:10.1038/ismej.2013.237.

Silman, N. J., M. A. Carver, and C. W. Jones. 1989. Physiology of Amidase Production by Methylophilus methylotrophus: Isolation of Hyperactive Strains Using Continuous Culture. Microbiology 135:3153–3164, doi:10.1099/00221287-135-11-3153.

Smith, H. L., and P. Waltman. 1995. The theory of the chemostat: Dynamics of microbial competition. Cambridge university press.

Sogin, M. L., H. G. Morrison, J. A. Huber, D. M. Welch, S. M. Huse, P. R. Neal, J. M. Arrieta, and G. J. Herndl. 2006. Microbial diversity in the deep sea and the underexplored “rare biosphere”. Proceedings of the National Academy of Sciences 103:12115–12120, doi:10.1073/pnas.0605127103.

Solorzano, L. 1969. Determination of ammonia in natural waters by the phenolhypochlorite method. L&O 14:799–801.

Sommer, U. 1981. The role of r-and K-selection in the succession of phytoplankton in Lake Constance. Acta Oecologica, Oecologia Generalis 2:327–342.

Steen, A. D., A. Crits-Christoph, P. Carini, K. M. DeAngelis, N. Fierer, K. G. Lloyd, and J. Cameron Thrash. 2019. High proportions of bacteria and archaea across most biomes remain uncultured. The ISME Journal 13:3126–3130, doi:10.1038/s41396-019-0484-y.

Teng, W., J. Kuang, Z. Luo, and W. Shu. 2017. Microbial Diversity and Community Assembly across Environmental Gradients in Acid Mine Drainage. Minerals 7:106.

Thingstad, T. F., and R. Lignell. 1997. Theoretical models for the control of bacterial growth rate, abundance, diversity and carbon demand. Aquatic Microbial Ecology 13:19–27, doi:10.3354/ame013019.

Tilman, D. 1980. Resources: A Graphical-Mechanistic Approach to Competition and Predation. The American Naturalist 116:362–393.

Tringe, S. G., C. von Mering, A. Kobayashi, A. A. Salamov, K. Chen, H. W. Chang, M. Podar, J. M. Short, E. J. Mathur, J. C. Detter, P. Bork, P. Hugenholtz, and E. M. Rubin. 2005. Comparative Metagenomics of Microbial Communities. Science 308:554–557.

Vallino, J. J. 2011. Differences and implications in biogeochemistry from maximizing entropy production locally versus globally. Earth Syst.Dynam. 2:69–85, doi:10.5194/esd-2-69-2011.

Vallino, J. J., C. K. Algar, N. F. González, and J. A. Huber. 2014. Use of Receding Horizon Optimal Control to Solve MaxEP-Based Biogeochemistry Problems. Pages 337–359 in R. C. Dewar, C. H. Lineweaver, R. K. Niven, and K. Regenauer-Lieb, editors. Beyond the Second Law - Entropy production and non-equilibrium systems. Springer Berlin Heidelberg, doi:10.1007/978-3-642-40154-1_18.

Vallino, J. J., and C. S. Hopkinson. 1998. Estimation of dispersion and characteristic mixing times in Plum Island Sound estuary. Estuarine, Coastal Shelf Sci. 46:333–350.

Vallino, J. J., and J. A. Huber. 2018. Using Maximum Entropy Production to Describe Microbial Biogeochemistry Over Time and Space in a Meromictic Pond. Frontiers in Environmental Science 6:1–22, doi:10.3389/fenvs.2018.00100.

Vallino, J. J., and I. Tsakalakis. 2020. Phytoplankton Temporal Strategies Increase Entropy Production in a Marine Food Web Model. Entropy 22:1–25, doi:10.3390/e22111249.

Violle, C., M.-L. Navas, D. Vile, E. Kazakou, C. Fortunel, I. Hummel, and E. Garnier. 2007. Let the concept of trait be functional! Oikos 116:882–892, doi:10.1111/j.0030-1299.2007.15559.x.

Wallenstein, M. D., and E. K. Hall. 2012. A trait-based framework for predicting when and where microbial adaptation to climate change will affect ecosystem functioning. Biogeochemistry 109:35–47, doi:10.1007/s10533-011-9641-8.

Weiss, R. 1970. The solubility of nitrogen, oxygen and argon in water and seawater. Deep-Sea Res. 17:721–735.

